# Rapamycin-PLGA microspheres induce autophagy and prevent senescence in chondrocytes and exhibit long *in vivo* residence

**DOI:** 10.1101/2020.04.06.027136

**Authors:** Kaamini M. Dhanabalan, Vishal K. Gupta, Rachit Agarwal

## Abstract

Osteoarthritis (OA) is a joint disease that results in progressive destruction of articular cartilage and the adjoining subchondral bone. The current treatment is focused on symptomatic relief due to the absence of disease-modifying drugs. The primary cells of the cartilage, chondrocytes, have limited regenerative capacity and when they undergo stress due to trauma or with aging, they senesce or become apoptotic. Autophagy, a cellular homeostasis mechanism has a protective role in OA during stress but gets downregulated in OA. Rapamycin, a potent immunomodulator, has shown promise in OA treatment by autophagy activation and is known to prevent senescence. However, its clinical translation for OA is hampered due to systemic toxicity as high and frequent doses are required. Hence, there is a need to develop suitable delivery carriers that can result in sustained and controlled release of the drug in the joint. In this study, we have fabricated rapamycin encapsulated poly (lactic-co-glycolic acid) (PLGA) based carriers that induced autophagy and prevented cellular senescence in human chondrocytes. The microparticle (MP) delivery system showed sustained release of drug for several weeks. Rapamycin-microparticles protected *in-vitro* cartilage mimics from degradation, allowing sustained production of sGAG, and demonstrated a prolonged senescence preventive effect *in vitro* under oxidative and genomic stress conditions. These microparticles also exhibited a long residence time of more than 19 days in the joint after intra-articular injections in murine knee joints. Such particulate systems are a promising candidate for intra-articular delivery of rapamycin for treatment of osteoarthritis.

**Statement of Significance:** Current OA treatment is symptomatic and does not change the disease progression. Many drugs have failed as they are rapidly cleared and are unable to maintain therapeutic concentration in the OA joint. Direct joint administration of drugs using sustained-release systems offer several advantages, which includes increased bioavailability, fewer off-target effects, and lower total drug cost. We have engineered a suitable drug carrier which provides a tunable drug release pattern. This study provides evidence that PLGA encapsulated rapamycin remained potent and prevented OA like changes in chondrocytes under genomic and oxidative stress. The particle formulation also had a longer residence time in the knee joint of mice which can be translated in clinics for intra-articular therapeutic injections for increased patient compliance.

**Graphical abstract:** 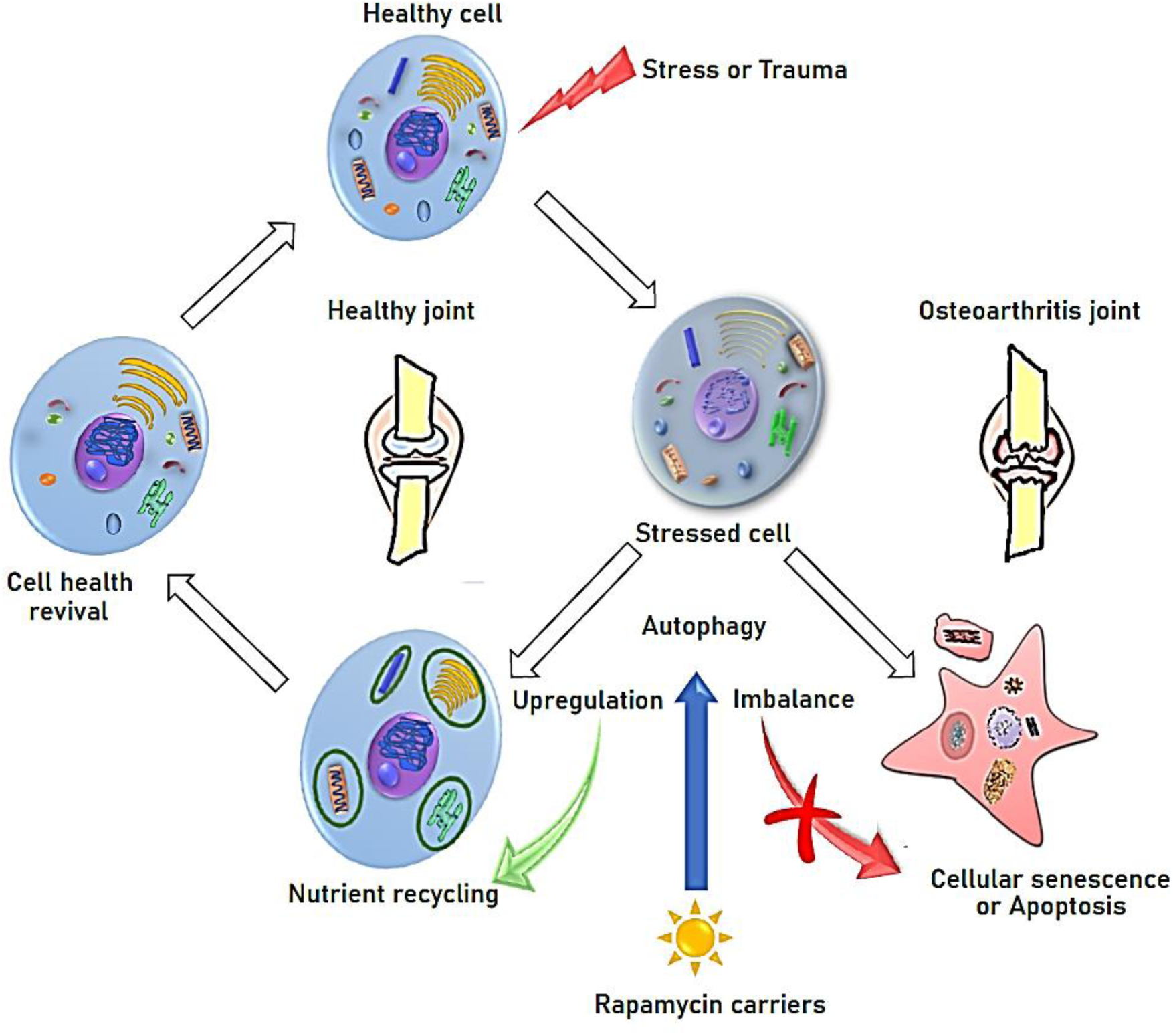

## Introduction

Osteoarthritis (OA) is a progressive degenerative joint disease that leads to poor quality of life majorly due to debilitation [1]. According to the WHO Global Burden of Disease Report 2018, it is estimated to affect 60-70% of the global population above the age of 65 and it is projected to increase as the global population continues to age. The current standard of care for OA revolves primarily around symptomatic relief and palliative therapies like giving anti-inflammatory drugs, pain killers and physiotherapy [2, 3]. Other treatments include invasive procedures like total knee replacements which are not affordable by the major part of the society or intra-articular injections with hyaluronic acid and corticosteroids which only show marginal improvements [2]. Importantly, no disease-modifying drug is available that can prevent, reverse or halt the progression of the disease. Several classes of drugs like cytokine receptor antagonists, anabolic growth factors, inhibitors of catabolic enzymes etc., have shown promise in preclinical trials but no approved product is in the market [2, 4, 5]. This is attributed to low bioavailability and rapid clearance of the injected drug from the target joint.

The key causes of OA are mechanical wear of human joints with age-related cartilage degeneration or trauma to joint. Cartilage and chondrocytes, the sole cellular residents of normal cartilage, have limited regenerative capacity [6, 7]. Articular cartilage damage and chondrocyte death occur with aging or after joint injury [1, 6, 8, 9]. Studies have suggested that with aging, there is an imbalance in cellular homeostasis which increases the risk of cartilage degeneration leading to OA [8, 10, 11]. In posttraumatic osteoarthritis, which occurs following an injury, the initial cartilage damage and stress prompts the chondrocytes to adopt a catabolic phenotype [12, 13].

Autophagy is a critical cellular homeostasis mechanism that supports normal cellular function and survival under stress-induced conditions such as nutrient deprivation and injury [14]. Autophagy is thus essential for maintaining cell and tissue homeostasis especially in knee joints given the limited blood supply and repeated mechanical stress on the joint [10, 11]. While the etiological cause of OA remains multifactorial, accumulating evidence has hinted towards an association between imbalanced autophagy in articular chondrocytes and the development of OA [10, 11, 15, 16]. Drugs that target autophagy induction have shown promise in OA and suggest that the regulation of autophagy may be a potential therapeutic strategy with which OA can be attenuated [17-20].

Another important aspect of OA disease progression is chondrocyte senescence (chondrosenescence). Senescence is a common molecular mechanism that potentially promotes both age-associated OA and trauma-induced OA [8, 21-23]. A recent study had shown that local clearance of senescent cells from the joint using UBX0101 (UBX), attenuated the development of post-traumatic osteoarthritis and created a pro-regenerative environment in mice [23]. This study has now further progressed to phase II (NCT04129944) clinical trials to evaluate the safety, tolerability and pharmacokinetics of UBX0101 in OA patients. Thus, targeting the senescent cells in the cartilage has emerged as an innovative strategy to stop the progression of OA. It has been reported that the chondrocytes in the cartilage are terminally differentiated and do not divide further [24]. Hence, it is important to prevent these limited number of cells from dying or becoming senescent.

Rapamycin, an FDA approved drug in the market for the past 20 years has been shown to prevent OA progression in mice models[17, 18, 25-27]. Rapamycin induces autophagy by inhibiting the mTOR pathway [18, 26]. When mice subjected to the destabilization of the medial menisci (DMM) were treated with rapamycin, a pharmacological activator of autophagy, chondrocyte cellularity was preserved and severe damage and degeneration was prevented in the articular cartilage [17, 18, 27]. Rapamycin is also known to be a senomorphic (senescence prevention) drug which can potentially prevent the cells under stress from turning into senescent phenotypes [28-31]. Thus, rapamycin-induced chondrocyte autophagy and senescence prevention can play a significant role in OA disease modification.

One of the major pitfalls associated with administering rapamycin as a free drug is its bioavailability at the target site. Large systemic doses are required to achieve therapeutic levels in the joints. Rapamycin exhibits systemic toxicity at such large doses leading to several side effects such as hypertriglyceridemia, hypertension, hypercholesterolemia, increased creatinine, constipation, abdominal pain, diarrhea [25]. To overcome these challenges, intra-articular (IA) administration of drugs are routinely used in the treatment of the joint disease and can target drugs directly to the affected tissues. The major advantage of this route of drug administration is that it minimizes the overall systemic exposure as well as allows a lower amount of drugs to be administered [32, 33]. Although intra-articular injections can ensure delivery of large doses in the joint, lymphatic clearance of drug can still rapidly reduce the drug concentration below the therapeutic levels. Drugs from the joint space are cleared at the rate of 0.04 mL/min causing very low residence time of low molecular weight drugs (∼1-4 h) such as rapamycin [32, 34]. As a result, frequent injections are required which can cause pain, patient non-compliance and potential infections.

In a study by Takayama *et al.*, intra-articular injection of rapamycin resulted in delayed degeneration of articular cartilage in the murine OA model. Since the drug was rapidly cleared, intra-articular injections were administered twice a week up to 8 weeks to achieve therapeutic efficacy [18]. Such a system is not translatable in clinics and may contribute to decreased patient compliance which reduces the overall efficacy of such treatments. Many drugs have been tested for OA but have failed to enter clinical practice owing to multiple weekly injections [2]. For instance, IL1-RA (Anakinra) therapy failed in clinical trials and its short half-life (2-4 h) is considered to be the primary reason for its failure [35]. Hence, there is a need for sustained drug release systems that can substantially increase the drug residence time to fully realize the potential of IA drug administration for OA treatment.

In order to overcome the limitation of poor drug retention in knee joints, the current focus has shifted towards using biomaterial-based microparticles, nanoparticles and hydrogels that can remain in the joint longer and release drug over time. Previous research and clinically available controlled-release depots (such as Lupron Depot^®^ and Risperdal Consta^®^) have shown that one can extend the residence time of the drug locally by encapsulating the drug in a polymer matrix, which slowly degrades in the biological medium releasing the drug [27, 36, 37]. Polymeric microparticles (MPs) are a feasible solution for controlled release of drugs due to their ability to encapsulate drugs with poor aqueous solubility while serving as controlled release systems which can potentially reduce the number of doses required to reach the desired therapeutic effect allowing for direct administration to the target site [13, 32, 34, 38-40]. Previous attempts to deliver rapamycin through such carriers have shown promise. In one study, gelatin hydrogels incorporating rapamycin–micelles were placed intra-articularly after surgery and were able to reduce the progression of experimental osteoarthritis [27] in a murine model. This suggests that extended release of rapamycin could help in OA disease alleviation.

Poly (D, L-lactic-co-glycolic acid) (PLGA)/ microspheres/nanoparticles are one of the most successful drug delivery systems (DDS) in labs and clinics. Because of their biocompatibility and biodegradability, they have been used in several biomedical applications such as drug delivery and tissue engineering [36]. PLGA encapsulated drugs like Lupron Depot^®^ & Vivitrol^®^ are successfully sold in the market and are widely used in clinics. Zilretta™ is a PLGA encapsulated triamcinolone acetonide formulation that has been widely used in clinics for reducing inflammation in OA patients and shows the feasibility of biomaterial-based delivery for OA. However, delivery of rapamycin for the treatment of OA through such clinically translatable materials like PLGA has not been explored yet.

Here, we present a robust drug carrier platform using PLGA, that can encapsulate active rapamycin and can be tuned to release the drug at varying rates for several days. The rapamycin encapsulated PLGA MPs stayed active, induced autophagy and prevented senescence under different stress conditions both in monolayer cultures as well as in 3D micro mass cultures. Administration of such MPs in the mice knee joint through intra-articular injections resulted in high residence time and colocalization with various joint tissues for up to 19 days.

## 2. Materials and Methods

### Mice

All animal experiments were approved by the Institutional Animal Ethics committee (IAEC) (CAF/Ethics/612/2018). The design, experimentation and analysis of animal experiments followed the IAEC guidelines. Mice were housed in a specific-pathogen-free animal facility at IISc Central Animal Facility. The animals were kept under constant temperature and humidity with automated 12 h light-dark cycles. Mice were fed standard laboratory food and water. Female wild-type (WT) mice of C57BL/J6 strain were used for intra-articular injection studies.

### Materials

Polymers and reagents Poly (D, L-lactic-co-glycolic) acid (PLGA, 50:50 and 65:35) of different molecular weights from 10,000 - 85,000 Da having carboxylic acid end groups were purchased from Akina (AP041) (West Lafayette, Indiana) and Sigma (719870, 719900, 719862) (St Louis, MO, USA). All other chemicals were of analytical grade and were used without further purification. Insulin/ transferrin /selenium (ITS) purchased from Gibco (Carlsbad, California, USA) were used for micro mass culture of the human chondrocyte cell line (C28/I2). Rapamycin (Sirolimus, 99.5% purity) was procured from Alfa *Ae*sar (J62473) (Ward Hill, Massachusetts, USA). Poly (vinyl alcohol) (PVA, 87∼89% hydrolysed, M_w_ 13,000 - 23,000 Da) was purchased from Sigma (363170) (St Louis, MO, USA). All other chemicals were of analytical grade obtained from Sigma Aldrich and used as received. **Antibodies used:** Rabbit Anti LC-3B polyclonal antibody (Abcam, Cambridge, United Kingdom, ab48394), Alexa Fluor 488 goat anti-rabbit IgG (H+L) (Abcam, Cambridge, United Kingdom A11008), Rabbit Polyclonal to *γ*H2Ax (Thermo Fischer, Waltham, Massachusetts, United States, PA5-28778).

## Methods

### 2.1 Synthesis of PLGA carriers with or without Rapamycin

Micro particles were synthesized using PLGA polymer of different molecular weights, using single emulsion method. Briefly, 100 mg PLGA was dissolved in 2 mL of Dichloromethane (DCM) with or without Rapamycin (1 mg) and homogenized in 10 mL of 1% PVA (aqueous phase) at 12000 rpm. This solution was added to 110 mL of 1% PVA (evaporation phase) and was allowed to stir continuously for 3-4 h to evaporate DCM completely. The solution was then centrifuged at 11,000 *g* and washed using deionized water twice to wash away the excess PVA. The microspheres were then re-suspended in deionized water and were rapidly frozen at -80° C followed by lyophilization.

### 2.2 Physiochemical characterisation of Microparticle formulation

#### 2.2.1 Encapsulation efficiency

To measure encapsulation efficiency, 10 mg of dried microspheres containing rapamycin were dissolved in 1 mL of Dimethyl Sulfoxide (DMSO) followed by absorbance measurement using a plate reader at 278 nm. The calibration curve for the quantification of Rapamycin and PLGA in DMSO was obtained. The encapsulation efficiency (**Table 1**) was calculated using the following equation:

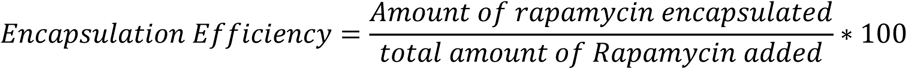

#### 2.2.2 Particle size

Particle size distributions were determined by dynamic laser light scattering (Nano ZS ZetaSizer (Malvern Instruments, Worcestershire, UK)). The PLGA microspheres (50 μg) were dispersed in deionized water (1 mL) and sonicated briefly before analysis. The volume mean diameter was used as a representative diameter.

#### 2.2.3 Scanning electron microscopy

For morphological examination and re-examining the particle size,10 µL of PLGA microparticles (100 µg/mL in distilled water) were dispersed on a metal stub and allowed to dry. The samples were then placed in a sputter coater for 45 s to produce a gold coating of 15 nm in thickness and rastered using an e-beam (15 KV) and viewed under a scanning electron microscope.

### *In vitro* release profile of rapamycin loaded micro particles

Rapamycin loaded micro particles (RMPs) (1 mg) were kept in 2 mL of 1x PBS solution (pH 7.4) at 37° C in a rotatory spinner inside the incubator. At various incubation times, the particles were pelleted at 11,000 *g* and the pellet was dissolved in 100 µL DMSO. The dissolved particles were quantified through plate reader instrument (Epoch™ 2 Microplate Spectrophotometer) with pre-determined parameters set to record the absorbance (278 nm). The amount of rapamycin released from each time point was extrapolated by using the rapamycin and PLGA standard curve. Blank micro particles were also subjected to the same as above. The experiment was carried out in triplicates.

### 2.4 Cell assays

#### Cell line and culture conditions

The human immortal chondrocyte cell line – C28/I2 (Merck, Kenilworth, New Jersey, United States; SCC043) were cultured in DMEM/F-12 medium (Invitrogen, Carlsbad, California, USA; 11320082) containing 1 mM sodium pyruvate, 10 mM of HEPES, 140 mM of glucose, supplemented with 10% fetal bovine serum (Sigma, F2442) and 1% penicillin/streptomycin (CC4007, Cell Clone, Delhi, India) in an atmosphere of 5% CO_2_ and 37° C. All the experiments in this study were based on this cell line.

#### 2.4.1 MTT assay

Before addition in the *in vitro* cell-based experiments, both empty and rapamycin-loaded MPs were sterilized by UV irradiation for 20 mins.

Cell’s metabolic activity was determined by 3- [4, 5-dimethylthiazol-2-yl]-2, 5-diphenyltetrazolium bromide (MTT) assay (Cayman chemicals; 1000936-5-500). This was done to determine the safe range of PLGA MPs concentrations that could be used for future experiments without causing a fall in the cell’s metabolic activity. Briefly, cells were plated at 15,000 cells per well in a 96 well plate and allowed to adhere overnight. Next day, PLGA microparticles without any drug were added in increasing concentrations (50 μg/mL, 100 μg/mL, and 500 μg/mL) and the cells were incubated for 24 h and 48 h in two independent experiments. After the incubation time, MTT reagent (10 µL) was added and the cells were incubated for 4 h, followed by crystal dissolving solution (CDS) to dissolve the formazan crystals. Absorbance readings were recorded at 570 nm.

#### 2.4.2 Live dead staining

Live dead staining was done using Calcein AM/Propidium Iodide (PI) staining. Briefly, 15,000 cells were plated in a 24 well plate the previous day and were allowed to adhere and grow overnight. Next day, rapamycin at different concentrations (10 nM, 1 μM, 100 μM) was added to the plated cells and incubated for 48 h. After the incubation time, the cells were thoroughly washed and Calcein AM/PI staining (final concentration 1 µg/mL) was done for 10-15 mins. The cells were then washed and imaged using IN Cell Analyzer 6000 (GE Life Sciences) in FITC (495/519 nm) and dsRed (558/583 nm) channels to evaluate for any cell death.

#### 2.4.3 Senescence induction assay

We subjected chondrocytes to two different stress mechanisms that are expected to be experienced by chondrocytes in OA. We treated chondrocytes to a genomic stress agent – BrdU (200 µM) (Bromo-4-deoxyUridine) which causes DNA damage, simulating aging-related OA. We also exposed the cells to oxidative stress agent H_2_O_2_ (200 µM) (Hydrogen peroxide) to simulate post-traumatic OA where there occurs a surge in ROS production upon stress to the cartilage [41]. Cells not treated with rapamycin/other stress agents received equal amounts of the vehicle. Colorimetric SA-β Gal activity was used to stain for senescent cells as previously described [42]. After exposing the cells to these stress agents for 48 h, cells were washed 3x with 1x PBS and then fixed with fixative solution containing 2% formaldehyde and 0.2% glutaraldehyde in 1x PBS for 10 min. Following fixation, cells were incubated in SA-β Gal staining solution (1 mg/mL 5-bromo-4-chloro-3-indolyl-beta-d-galactopyranoside (X-Gal), 1x citric acid/sodium phosphate buffer (pH 6.0), 5 mM potassium ferricyanide, 5 mM potassium ferrocyanide, 150 mM NaCl, and 2 mM MgCl_2_) at 37° C in a non-CO_2_ incubator for 16–18 h. The enzymatic reaction was stopped by washing the cells three times with 1x PBS. Cells were counterstained with DAPI (300 nM) solution for 10 mins and then rinsed with 1x PBS. Five random images per well were taken in bright field channel followed by the respective fields in the DAPI channel. These images were analysed using ImageJ software using a pre-written macros program to automatically count the senescent cells in bright field channel and the DAPI channel was used to identify the total number of cells. From these, the percentage of senescent cells was calculated.

#### 2.4.4 Rapamycin MPs treatment in senescence induction assays (Monolayer cell culture)

For the experiment, the chondrocytes (C28/I2) were plated in a 24 well plate with 7,500 cells per well in the untreated group and 15,000 cells per well for other treatment groups to maintain cells under sub confluency until the end of the experiment. The cells were treated with either BrdU/H_2_O_2_ (200 μM) and co-treated with Rapamycin loaded MPs (final concentration 1 μM, 200 nM and 50 nM) through a tissue culture plate inserts (pore size 0.1 µm) (VWR^®^ 10769-220) assay to prevent uptake of MPs in the chondrocytes which interferes with downstream SA-β Gal staining. After 48 h of incubation, the cells were fixed using the fixative solution containing 2% formaldehyde and 0.2% glutaraldehyde in 1x PBS followed by SA-β Gal staining for identifying senescent cells as well as DAPI to stain the nucleus.

In order to automatically score the senescent cells, we developed a custom-built macros algorithm to score the senescent cells based on their size and the intensity of SA-β Gal staining. Brightfield images were taken from each treatment groups (vehicle (DMSO) treated cells, BrdU/H_2_O_2_ treated cells, BrdU/H_2_O_2_ and rapamycin treated cells etc). The macros algorithm was run to count the total number of senescent cells.

To count the total number of cells, we used DAPI to stain the nucleus. The images that were obtained in the DAPI channel for the corresponding images in brightfield were used to count the total number of cells in that image area. A customised macros algorithm was used to automatically count the nucleus, indicative of cell count.

#### 2.4.5 Rapamycin MPs treatment in senescence induction assays (Micro mass culture)

We generated micro masses as described previously [43]. Briefly, the cells were seeded as a 15 μL suspension in growth media in a 24 well plate at a density of 2 x 10^7^ cells/ mL. The cells were allowed to adhere to the well plate for 3 h after which growth media was added. After 24 h, the growth media was changed to differentiation media containing various supplements like Insulin/Transferrin/Selenium, TGF-β (10 ng/mL), and ascorbic acid. They were then divided into their respective treatment groups and were treated with BrdU (600 μM) /H_2_O_2_ (100 μM) and co-treated with rapamycin (1 μM)/rapamycin loaded MPs (Final concentration 1 μM, 200 nM and 50 nM). After 48 h of incubation, the cells were fixed using 4% formaldehyde followed by Alcian blue (Sigma-Aldrich, A5268) staining at pH<1 to stain the sulphated glycosaminoglycans(sGAG) overnight. The next day the micro masses were washed to remove any non-specific stains followed by Alcian blue stain extraction using Guanidine HCl (Sigma Aldrich, SRE0066). The extracted Guanidine HCl’s absorbance was read at 630 nm using a plate reader (Epoch™ 2 Microplate Spectrophotometer) to quantify the proteoglycans present in the micro masses after various treatments. Similar experiments were carried out with incubation time of up to 8 days with BrdU/H_2_O_2_ and BrdU/H_2_O_2_ along with rapamycin MPs for 8 days. Media was changed every two days once and were replenished with BrdU/H_2_O_2_ along with rapamycin MPs in the respective treatment groups.

#### 2.4.6 Immunocytochemistry to detect *γ*H2Ax

To check the ability of rapamycin MPs in preventing DNA damage under genomic stress conditions using BrdU (200 μM), we treated sub-confluently plated chondrocytes with BrdU (200 μM) and co-treated other groups with BrdU (200 μM) and rapamycin (1 μM, 200 nM and 50 nM) for 48 h. After the incubation time, the cells were fixed with 4% PFA for 15 minutes at room temperature followed by permeabilization using 0.05% Triton X-100. The cells were then blocked using 5% non-fat dry milk and stained using Rabbit Anti *γ*H2Ax polyclonal antibody (0.21 µg/mL) in the nucleus followed by incubation with Alexa Fluor 488 goat anti-rabbit IgG (H+L) (0.42 μg/mL) for 1 h and nuclear staining using DAPI (300 nM) for 10 mins. These cells were then visualised using FITC (495/519 nm) and DAPI (358/451 nm) channel using IN Cell Analyzer 6000 (GE Life Sciences) at 40x magnification.

#### 2.4.7 Immunocytochemistry to detect LC3B

Here the chondrocytes (C28/I2) were plated in a 24 well plate with treatments of with and without rapamycin/rapamycin MPs (Final concentration 1 μM, 200 nM and 50 nM). After 6 h of treatment, the cells were fixed using 4% PFA followed by permeabilization using 0.05% Triton X-100. The cells were then blocked using 5% non-fat dry milk and incubated overnight at 4° C with Rabbit Anti LC-3B polyclonal antibody (0.5 μg/mL). Next day the cells were washed and incubated with Alexa Fluor 488 goat anti-rabbit IgG (H+L) (1 μg/mL) for 1 h followed by nuclear staining using DAPI (300 nM) for 10 mins. These cells were then visualised using FITC (495/519 nm) and DAPI (358/451 nm) channel using IN Cell Analyzer 6000 (GE Life Sciences) at 40x magnification.

### 2.5 *In vivo* residence time of PLGA MPs in mouse knee joints

PLGA particles of M_w_ 75 KDa-85 KDa were synthesised by single emulsion method as described earlier, with Cy5 (50 µg) dye added to the organic phase during synthesis. The particles were lyophilised, sterilised by UV irradiation before making MP formulation in 1x PBS.

A sterile suspension of 20 mg/mL particles was prepared and 10 µL from this formulation was injected into mice knee joint space on Day 0. Two mice were sacrificed on day 1, 3, 7, 14 and 19 for knee joints dissection and joint components were separated in a 24 well plate followed by fixation in 4% formaldehyde and stained using 600 nM of DAPI for 10 minutes. The joints were then washed with 1x PBS and then imaged using IN Cell Analyzer 6000 (GE Life Sciences) at 20x magnification to check for Cy5 particles localisation. The joints were imaged in DAPI (358/451 nm) channel and Cy5 (678/694 nm) channel with the femur, tibia, meniscus and tibia separately.

## Statistical analysis

Data is presented as mean ± standard deviation. Differences between groups were analyzed by the Student’s *t*-test, One-way ANOVA of variance or Kruskal Wallis test depending on the number of variables, with *p <*0.05 considered significant.

## 3. Results

### 3.1 PLGA microspheres fabrication and characterization

For longer residence in the joint, size is an important parameter for the effectiveness of the treatment, as it directly influences the retention of the drug in the joint. We have previously shown that about 900 nm - 1 µm particles showed increased residence time in rodent knee joints [44]. Based on this, we synthesized 1 µm particles by adopting the previously described protocol [45]. **Figure 1a** shows PLGA MPs containing Cy5 fluorescent dye. Scanning electron microscopy (SEM) showed spherical shape and smooth surface with an average diameter of about 1 µm, which is in agreement with the results from light scattering measurements (**Figure 1b**). Dynamic light scattering for size analysis revealed an average diameter of 1026 ± 153 nm (**Figure 1c**).

**Figure 1:**
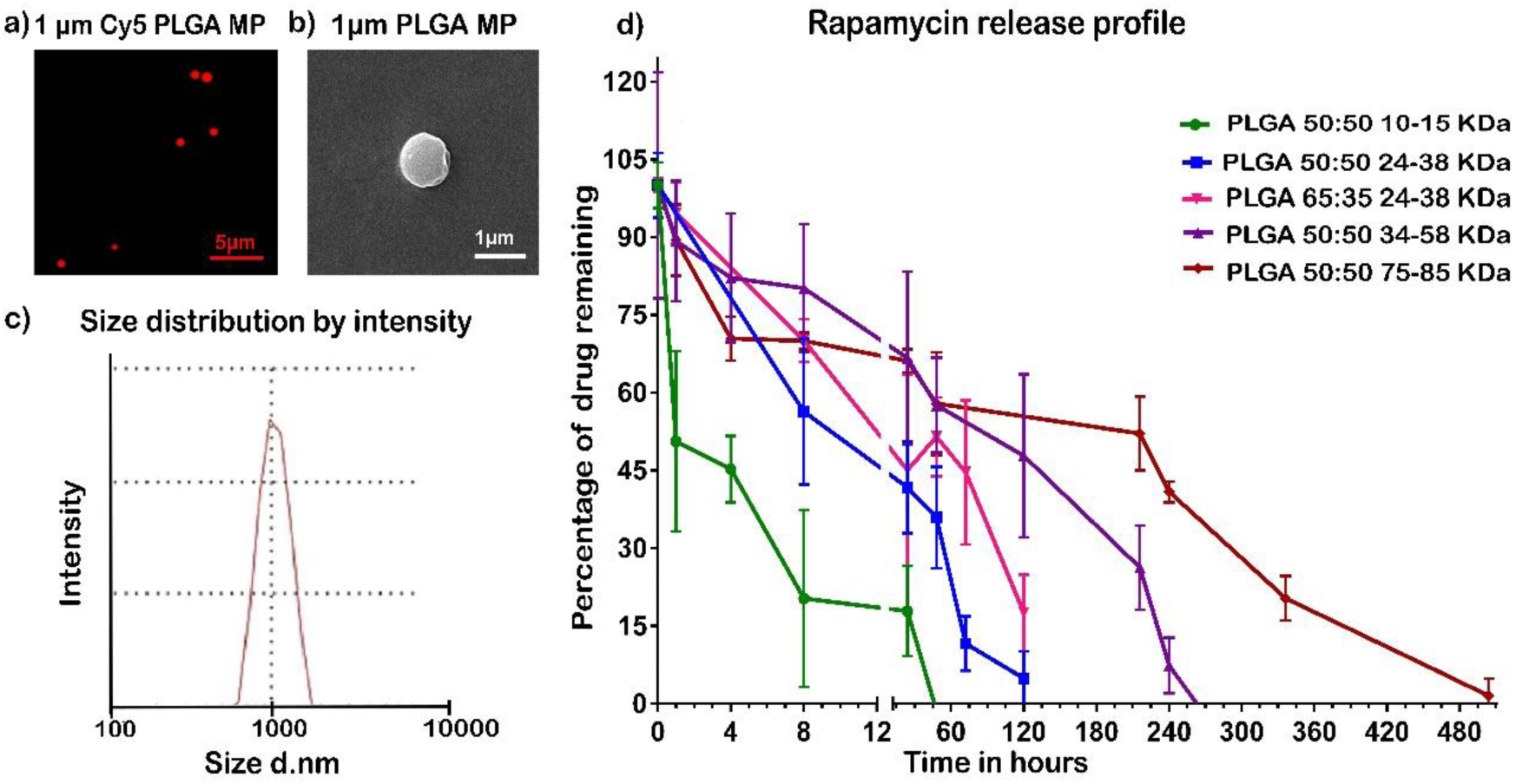
Characterization of PLGA MPs and drug release. (a) Fluorescence microscopy image of Cy5 loaded PLGA particles, Scale Bar 5 μm (b) Scanning Electron Micrograph of PLGA particle, Scale Bar 1 μm (c) Size distribution of 1 μm particles as measured by Dynamic Light Scattering (d) Quantification of *in vitro* release profiles of rapamycin from PLGA particles synthesized from different molecular weight polymer.

In order to vary the drug release profile, from the prepared MPs, different molecular weight polymers were used to encapsulate rapamycin. The size of the MPs and their encapsulation efficiency is given in Supplementary data **Table S1**. Particles prepared from different molecular weight polymers showed similar size range of about 1 µm and encapsulation efficiencies were varied from 32-65%.

### 3.2 PLGA microparticles exhibited controlled and tunable release of rapamycin

Rapamycin MPs which were kept in PBS in a roto spin were pelleted down at different time points and the pellet was dissolved in DMSO to release the remaining rapamycin. Three replicates were used at each time point. All particle formulations showed a sustained release of rapamycin for various durations (**Figure 1d**). Release profiles of PLGA (different molecular weights) showed that rapamycin was gradually released from the higher molecular weight PLGA MPs (75-85 KDa) until 21 days while the lower molecular weight PLGA MPs (10-15 KDa) released the drug completely within 48 h. We also tried using PLGA with 65:35 ratio of PLA and PGA to check if the release could slow down owing to higher crystalline nature of PLA. It has been shown in literature that PLA is highly crystalline in nature and exhibits hydrophobic property [46, 47]. Hence PLGA polymers containing higher ratios of PLA may contribute to slower release of the drug when compared to 50:50 ratios of PLA and PGA. Use of 65:35 ratio polymer allowed us to slightly increase the drug release time but effect of varying molecular weight was more prominent. This shows that PLGA particles provide a good platform to vary the release of the drug and can be used for sustained release of drug for several weeks. The 10-15 KDa M_w_ PLGA MP had the fastest rapamycin release rate (complete release within 48 h). Since for *in vitro* experiments, cell culture media is to be changed every 2-3 days, we used 10-15 KDa M_w_ PLGA MP for all *in vitro* experiments.

### 3.3 PLGA MPs are cytocompatible

In order to assess the cytocompatibility of our particle formulations, the metabolic activity of the cells in the presence of MPs was tested. We subjected the human chondrocytes (C28/I2) to different concentrations of PLGA particles for 24 and 48 h (Supplementary **Figure S1a and b**). Post incubation time, it was observed that the cell’s metabolic activity started to reduce only when the concentration reached 500 µg/mL after 48 h. Therefore, for all future experiments, we have used less than 100 μg/mL of MPs.

### 3.3 Live dead staining to determine the cytotoxicity of Rapamycin as a free drug

We next determined the cytotoxic effects of varying rapamycin concentrations on human chondrocytes. Live/ dead staining was performed after rapamycin treatment for 48 h to see the cytotoxic effect. Rapamycin did not cause cell death (Supplementary **Figure S1c-f**), even up to 100 µM concentration evidenced by very few PI stained cells in each panel of the image.

### 3.5 Rapamycin MPs induced autophagy in chondrocytes

LC3B is a common marker used to identify autophagy induction [48]. During autophagy signalling, the ubiquitously present LC3B protein gets converted from LC3B-I isoform to LC3B-II isoform by phosphatidylethanolamine conjugation and gets inserted into autophagosome membrane [48-50]. Thus, a diffuse signal from the cytoplasm and nucleus gets converted into a punctate signal visualised by immunocytochemical staining [49, 50]. When chondrocytes were treated with rapamycin MPs (equivalent rapamycin dose of 1 µM, 200 nM or 50 nM) (**Figure 2d and Supplementary data Figure S2c, d respectively**) for 48 h, LC3BII was seen as punctate signals when compared to the untreated cells (**Figure 2a**) which showed diffused signals. The results from rapamycin MPs were comparable to that of the free rapamycin treated groups (**Figure 2b, and supplementary data Figure S2a, b respectively**). This experiment successfully showed that rapamycin was present in its active form in the microparticle formulation and induced autophagy in chondrocytes in a dose dependent manner.

**Figure 2:**
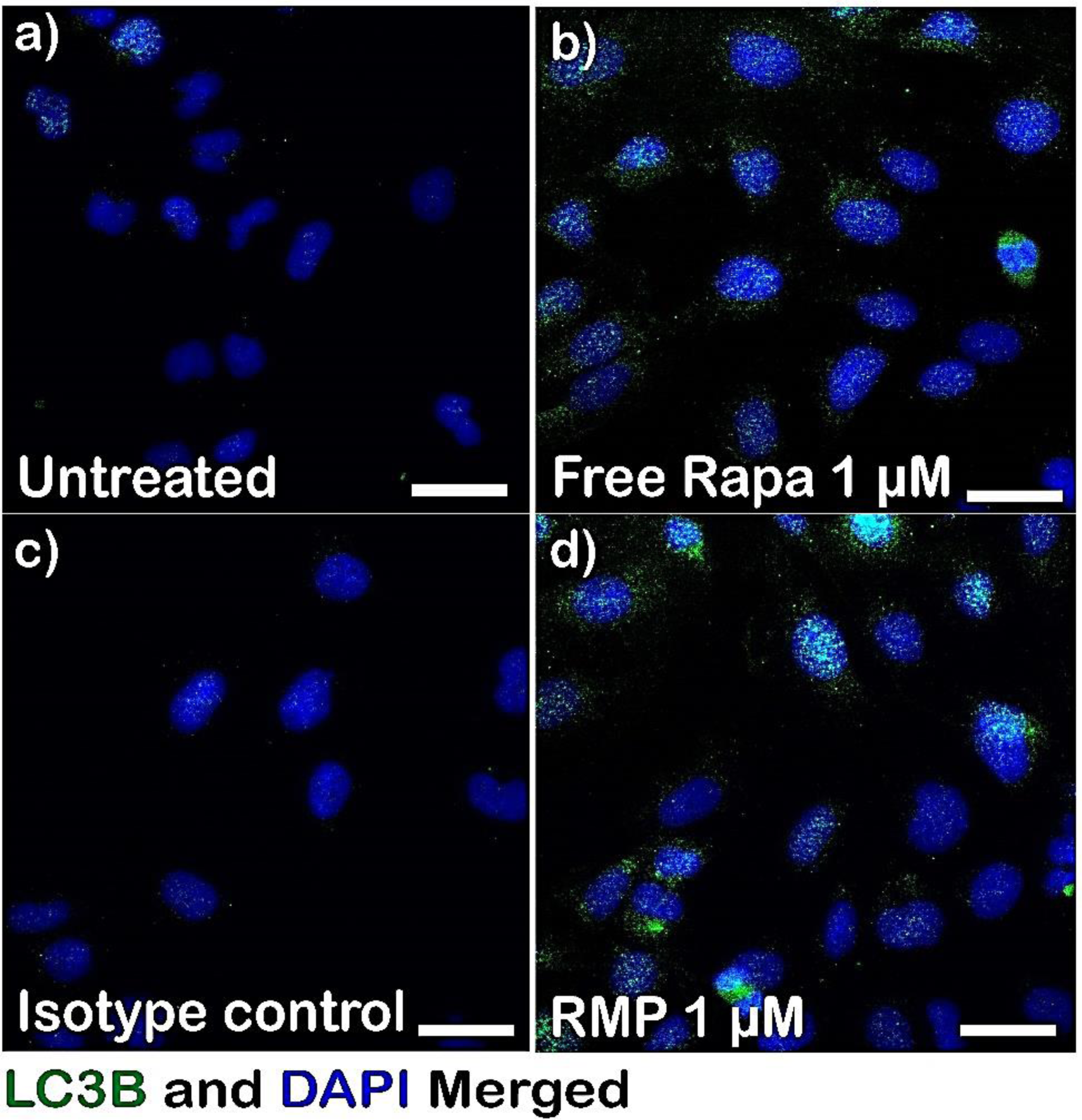
Rapamycin MPs induces autophagy in chondrocytes. Fluorescence microscopy images of C28/I2 cells stained with DAPI and LC3B primary antibody after treatment with (a) no rapamycin, (b) free rapamycin (1 µM) (c) Isotype control (d) Rapamycin MPs (1 µM rapamycin). Scale bar 20 μm.

### 3.6 Rapamycin prevented senescence in cells that were under genotoxic and oxidative stress (monolayer culture)

To mimic the cells under stress as is shown to be present in OA, we subjected the chondrocytes to two different stress conditions: genotoxic stress (BrdU) (simulating DNA damage like condition) and oxidative stress (H_2_O_2_) (simulating inflammation) [41, 51, 52] along with a co-treatment of rapamycin MPs at two different doses 200 nM and 50 nM. At the end of the incubation timepoints, the cells were fixed and stained for SA-β-Gal and DAPI. The images were scored by a custom-built automated macros algorithm in ImageJ for senescent cells as well as for total cells stained using DAPI.

When subjected to 200 µM of genotoxic stress agent (BrdU) about 55% of the cells became senescent while the rapamycin MPs co-treated groups – 200 nM and 50 nM of rapamycin, significantly brought down the percentage of senescent cells to 12.6% and 15% respectively (**Figure 3a, b and e**). These were comparable to that of the free rapamycin treated groups which brought down the senescent cells to 14%.

**Figure 3:**
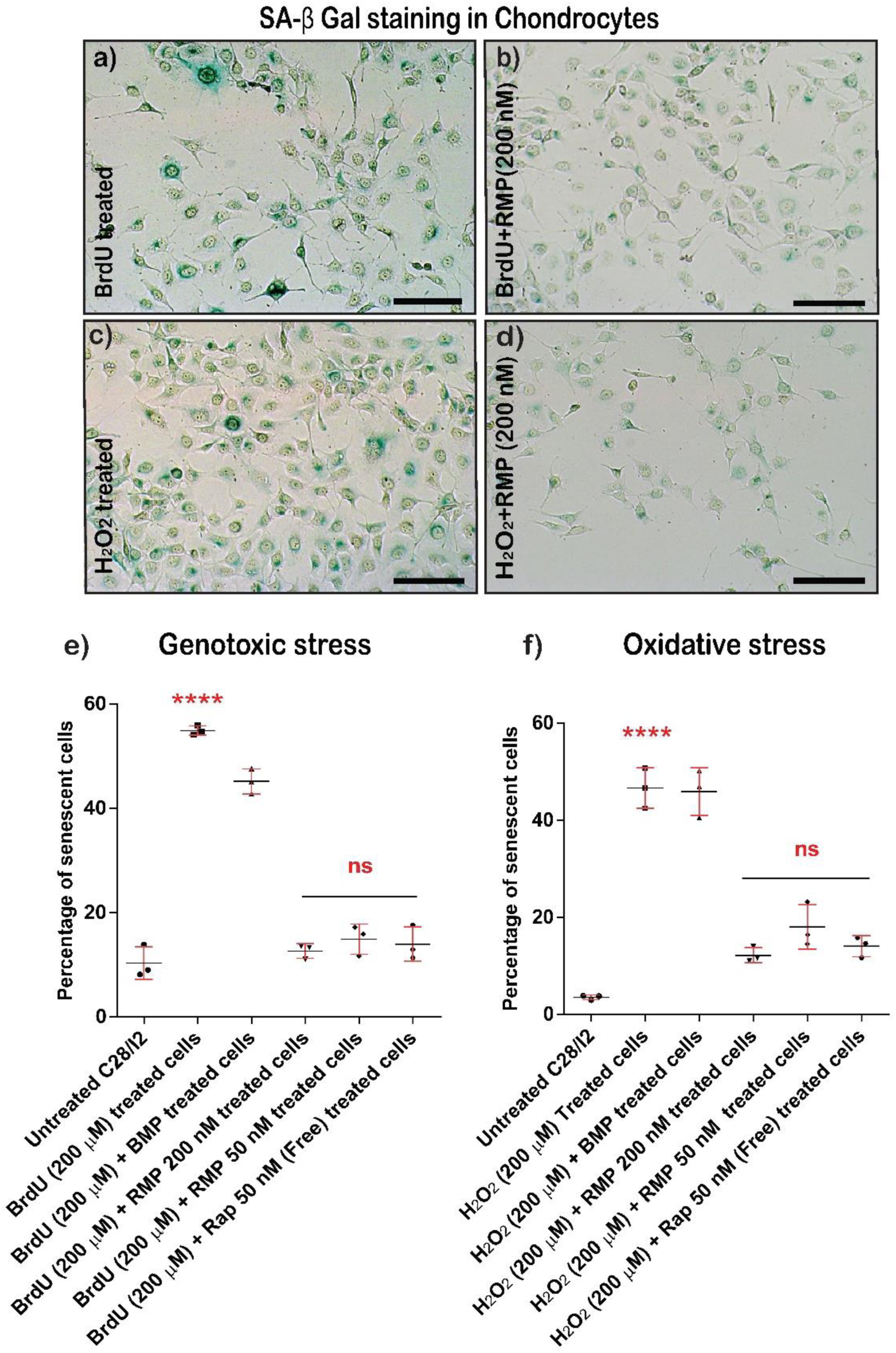
Rapamycin MPs prevent senescence in chondrocyte (C28/I2) culture. Brightfield images of C28/I2 cells stained with SA-β Gal exposed to (a) genotoxic (BrdU) stress (b) genotoxic (BrdU) stress along with Rapamycin MPs treatment (c) oxidative (H_2_O_2_) stress (d) oxidative (H_2_O_2_) stress along with Rapamycin MPs treatment. Scale bar 200 µm. Percentage of senescent cells under (e) genotoxic (BrdU) stress condition, (n = 3 per group) (f) oxidative (H_2_O_2_) stress condition (n = 3 per group). Data in graphs represent the mean ± s.d. and *p* values were determined by Kruskal Wallis test and Tukey’s post hoc tests using GraphPad Prism Software. For **e** and **f**, \*\*\*\**p* < 0.0001 for all rapamycin, RMP treated and untreated groups vs BrdU/ H_2_O_2_ treated group. *p*-value < 0.05 was considered significant. BMP-Blank Micro Particles, RMP-Rapamycin loaded Microparticles, ns-non-significant.

We then performed a similar experiment using an oxidative stress agent (H_2_O_2_ 200 µM) (simulating inflammation) and co-treated with rapamycin MPs (200 nM and 50 nM) for 48 h. H_2_O_2_ treated group had 46.6% of senescent cells while rapamycin MPs treated groups – 200 nM and 50 nM significantly brought the senescent cells down to 11.6% and 18% respectively (**Figure 3c, d and f**). These were again comparable to the free rapamycin treated groups which only had 14% senescent cells.

These data suggest that the rapamycin MPs were potent and active in the encapsulated form and were able to prevent the chondrocytes from going into senescence on par with free rapamycin ability.

### 3.7 Rapamycin MPs prevented the formation of heterochromatin nuclei in cells under genotoxic stress

Articular chondrocytes in joints affected by OA exhibit a senescent phenotype that is partly induced by the gradual accumulation of genomic DNA damage [41, 53]. One of the hallmarks of DNA damage repair mechanism is the formation of heterochromatin condensation foci [54]. The heterochromatin condensations contain phosphorylated Histone A protein (*γ*H2Ax) hence it is a widely used senescent marker [53, 54] which detects DNA damage response in the nucleus. Data suggests that strategies that result in the reduction of genomic DNA damage might help to prevent joint damage in OA [41, 52, 53, 55].

With our previous experiments, we have provided evidence that genotoxic stress agent-BrdU induces senescence in chondrocytes. This senescent like phenotype in chondrocytes *in vitro* exhibited increased SA-β Gal activity. We next wanted to test whether rapamycin MPs can prevent DNA damage from occurring in chondrocytes exposed to BrdU. The immunocytochemistry staining for (*γ*H2Ax) in the various treated groups show that rapamycin microparticle formulation was able to prevent DNA damage in a dose dependent manner, which is evident from the reduced number of (*γ*H2Ax) foci (**Figure 4c and supplementary data Figure S3a and b**) in the rapamycin MPs treated groups.

**Figure 4:**
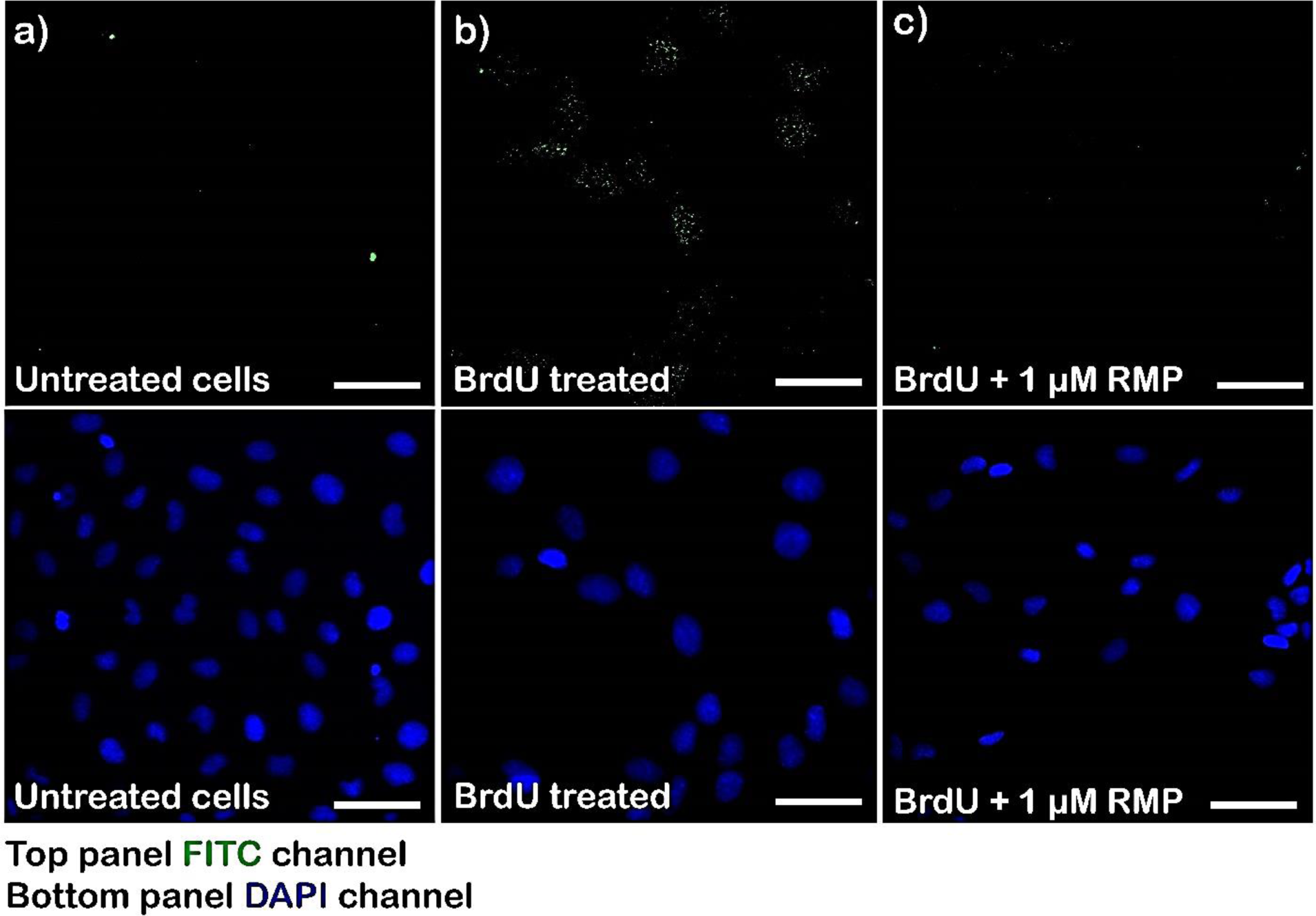
Rapamycin MPs reduces the DNA damage in cells under genotoxic stress: Fluorescence microscopy images of C28/I2 cells stained with DAPI and γH2Ax antibody after treatment with (a) no treatment (b) BrdU (200 µM) treated (c) BrdU (200 µM) along with Rapamycin MPs (1 µM rapamycin). Top Panel – FITC channel, Bottom panel – same field in DAPI channel. Scale bar 20 μm.

### 3.8 Rapamycin helps to sustain sGAG production in micro mass cultures under stress

When the chondrocytes are seeded at a high density in the presence of growth factors like TGF-β (10 ng/mL), they release extra-cellular matrix components like Collagen2A1 and Aggrecan [56]. The chondrocytes along with the secreted ECM and added growth factors together behave like a cartilage phenotype. These 3D cultures referred to as the micro mass cultures, are considered as *in vitro* mimic of cartilage and are widely used in the field of cartilage research [43, 56].

We used these micro mass cultures, derived from human chondrocyte cell line C28/I2 to evaluate whether free rapamycin and rapamycin MPs can sustain the production of sulphated glycosaminoglycans (sGAG) under different stress conditions both for short and longer duration of stress. The micro masses were treated with two different stress conditions similar to monolayer cultures along with co-treatment of rapamycin MPs at three different doses 1 µM, 200 nM and 50 nM. sGAG production was measured using Alcian blue staining as described [43, 56]. When exposed to 600 µM of BrdU for 48 h, the absorbance values dropped down to almost two folds while the 1 µM equivalent rapamycin delivery through rapamycin MPs had absorbance values similar to untreated samples indicating that the cells were able to sustain the sGAG production. This was comparable with free rapamycin group (**Figure 5a**). The lower doses of rapamycin MPs – 200 nM and 50 nM showed some improvements in sGAG content however were not significantly different from the BrdU treated group.

**Figure 5:**
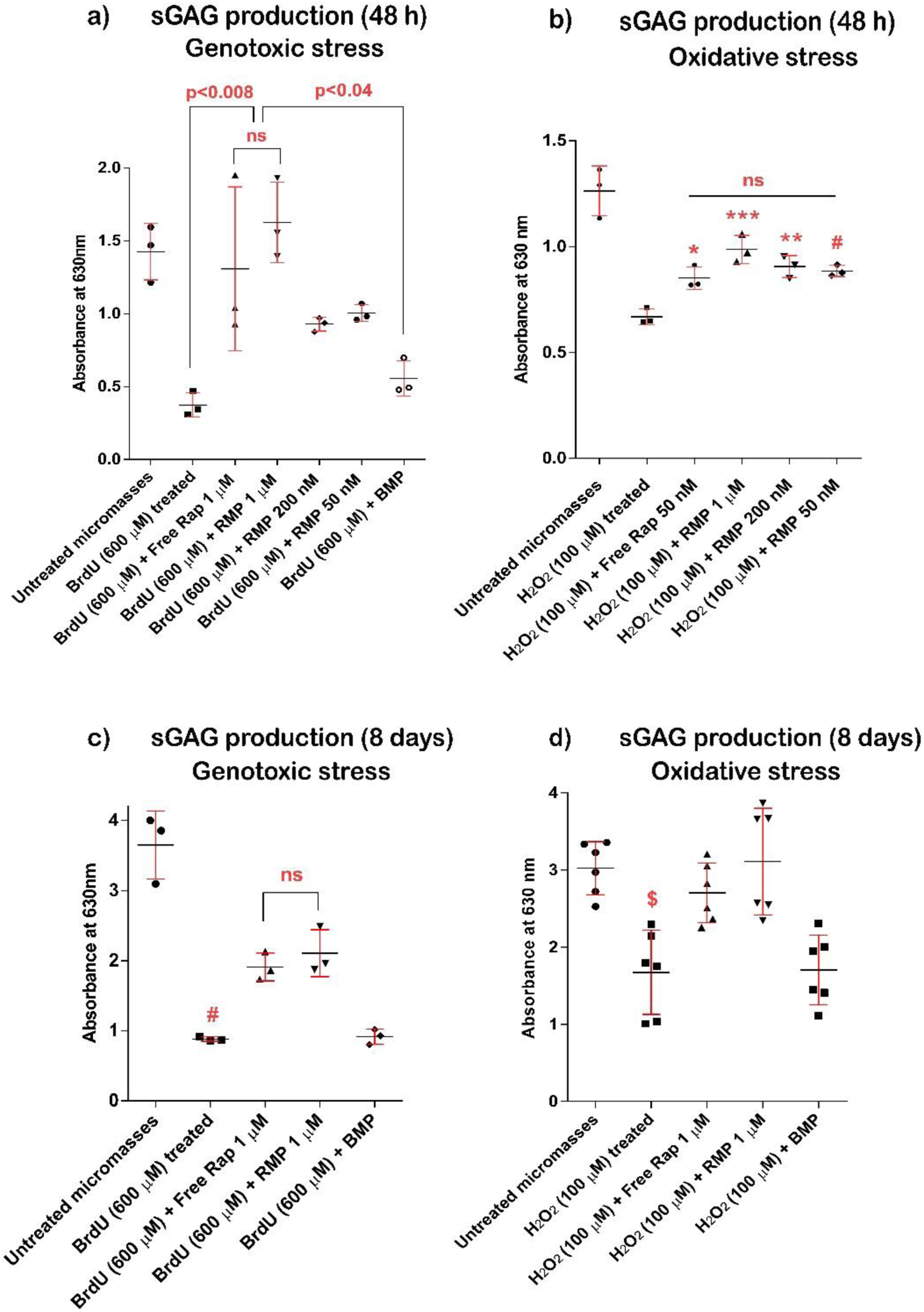
Rapamycin MPs prevent loss of sGAG in micro mass cultures exposed to genotoxic and oxidative stresses. sGAG production from C28/I2 micro mass culture after treatment with various particle and drug formulations under (a) genotoxic (BrdU) stress for 48 h (n = 3), (b) oxidative (H_2_O_2_) stress for 48 h (n = 3), (c) genotoxic (BrdU) stress for 8 days (n = 3) and (d) oxidative (H_2_O_2_) stress for 8 days (n = 6). Data in graphs represent the mean ± s.d. and *p* values were determined by one-way analysis of variance (ANOVA) and Tukey’s post hoc tests using GraphPad prism Software *p*-value < 0.05 was considered significant. For **b**, * indicates *p*=0.0458, \*\*\**p*=0.0008, \*\**p*=0.0082 and #*p*=0.0159. For **c**, # indicates *p*<0.0085 compared to untreated and all other rapamycin treated groups. For **d**, $ indicates *p*<0.0115 compared to untreated and all other rapamycin treated groups. BMP - Blank Micro Particles, RMP - Rapamycin loaded Microparticles, ns – non-significant.

Similar experiments were carried out using oxidative stress agent (100 µM H_2_O_2_) along with a co-treatment of rapamycin MPs at three different doses 1 µM, 200 nM and 50 nM. As evident from **Figure 5b**, the H_2_O_2_ treated micro masses exhibited much lower levels of sGAG content compared to untreated. All free and microparticle based rapamycin delivery significantly enhanced sGAG production compared to H_2_O_2_ treated micro masses. There was no significant difference between any of the rapamycin treatment groups however, all the groups were significantly lower than the untreated micro masses. Thus, both free rapamycin and rapamycin MPs were able to demonstrate intermediate sGAG production in these cartilage mimics at different doses for up to 48 h.

We next evaluated whether the rapamycin MPs will be able to sustain sGAG production in chondrocytes under a longer duration of stress (8 days). So, we performed similar experiments using BrdU (600 µM) and H_2_O_2_ (100 µM) along with co-treatment groups containing rapamycin MPs at 1 µM dosage for a duration of 8 days. On the 9^th^ day, the micro masses were stained for Alcian blue to evaluate the sGAG content in different treatment groups.

In the BrdU (600 µM) treated group, the absorbance values dropped to almost four folds at 630 nm while the rapamycin MPs treated groups restored the absorbance values which were comparable to the free rapamycin treated groups (**Figure 5c**). Similarly, after H_2_O_2_ treatment, micro masses had almost two-fold lower readings compared to the untreated micro masses at 630 nm. The rapamycin MPs treated group significantly increased the absorbance readings compared to the H_2_O_2_ treatment and were comparable to the untreated group (**Figure 5d**). These data are conclusive evidence of rapamycin MPs ability to sustain sGAG production by the chondrocytes under different stress conditions for long durations.

### 3.8 Rapamycin helps to sustain GAG production in micro mass cultures pre exposed to oxidative stress

We next wanted to evaluate whether rapamycin MPs will be able to sustain sGAG production even in chondrocytes that were pre exposed to oxidative stress conditions before treatment with rapamycin. For this study, we initially exposed the micro mass cultures to 100 µM of H_2_O_2_ for two days following which media was changed with treatment groups containing H_2_O_2_ as well as rapamycin MPs (1 µM) / free rapamycin (1 µM). The experiment was carried on for another 6 days with media change and fresh components added once every two days. On the 9^th^ day, the micro masses were stained for Alcian blue followed by guanidine HCl extraction and absorbance readings at 630 nm. The results (**Figure 6**) demonstrated that the rapamycin MPs had significantly higher (*p*=0.0305) sGAG production compared to H_2_O_2_ alone even in cultures pre exposed to oxidative stress. The sGAG production was similar to untreated when the micro masses were treated with free rapamycin (1 µM) while intermediate improvements were observed when treated with rapamycin microparticles. This could be attributed to the immediate availability of rapamycin in free drug group while in case of rapamycin MPs, there could be a delay in the release of sufficient rapamycin to protect cells that were already under the oxidative for two days.

**Figure 6:**
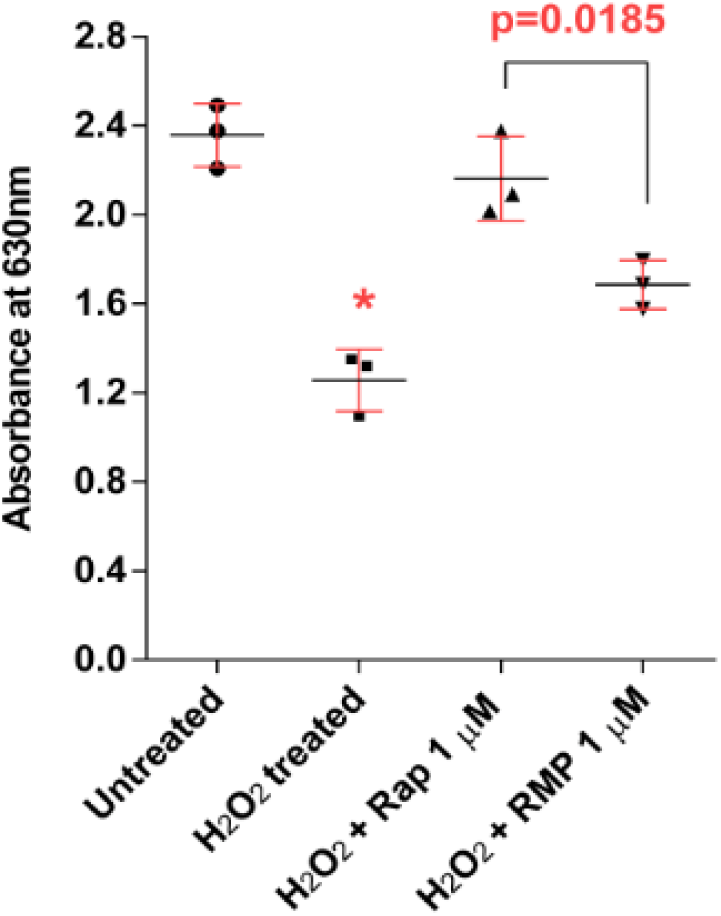
Rapamycin MPs prevent loss of sGAG production in micro mass cultures pre exposed to oxidative stress. sGAG production in C28/I2 micro mass cultures pre-exposed to oxidative stress (100 µM of H_2_O_2_) for 48 h followed by co-treatment with H_2_O_2_ and rapamycin formulations for 6 days. Data in graphs represent the mean ± s.d. (*n* = 3 per group) and *p* values were determined by one-way analysis of variance (ANOVA) and Tukey’s post hoc tests using GraphPad Prism software. *p*-value < 0.05 was considered significant. \**p*<0.04 against all other groups. BMP - Blank Micro Particles, RMP - Rapamycin loaded Microparticles, ns – non-significant.

### 3.10 High molecular weight PLGA MPs had a longer residence time in the mice knee joints

Results from the previous experiments showed that drug released from rapamycin MPs was potent and active. Rapamycin MPs successfully induced autophagy and prevented senescence in chondrocytes in *in vitro* conditions. In order to evaluate the robustness of this formulation in *in vivo* system, we studied the retention time of 1 µm particles in articular joint space of mice after intra-articular delivery. We used the PLGA MPs that had the higher M_w_ (75-85 KDa), as from the release studies it was evident that they slowly degraded over several weeks. Such higher M_w_ MPs can serve as a suitable candidate for potential *in vivo* treatments and hence we used them to check for residence time, localisation of the particles within the tissues in the joint. The animals were also monitored and assessed for any gross inflammation or infection post intra-articular injection of PLGA MPs. When MPs were injected into mice knee joints via intra-articular injection, the Cy5 loaded MPs were found to localise with femoral, tibial, synovial and meniscal surfaces at all time points tested (**Figure 7**). We found a gradual decrease in the number of particles in joint with time. Even at 19 days after injection, the particles were localized with various sections of the joint. Given the long residence time of these MPs and a gradual decrease in the signal, it is expected that the particles are degrading resulting in encapsulated Cy5 dye being released and cleared from the joint. We also observed that rapamycin showed complete release from these particles in 21 days (**Figure 1d**). However, macrophage uptake by macrophages may also be contributing to this clearance. The animals showed no macroscopic signs of swelling, infection or inflammation or any disability of the joint. This shows that the particles are safe to inject and have long residence time.

## DISCUSSION

Research has shown that OA can occur due to an imbalance between the anabolic and catabolic activity of chondrocytes [10, 16] and due to chondrosenescence [21, 22]. Defective autophagy is one such cause of OA which is closely implicated in the aging process and after trauma to the joints [10, 11, 16, 26]. Both, human OA, as well as surgically induced OA in mice, are associated with a reduction and loss of autophagy [10-12, 15, 16, 26]. It has been shown that autophagy descends to a downward spiral after its peak during the early stage of OA, which results in the aggravation of OA [10, 11, 15, 16, 20]. Recent reports show that activation of autophagy significantly delays cartilage degradation and can pave way for OA disease amelioration [11, 17, 19]. Rapamycin, a clinically used potent immunomodulator [25] and macrolide antibiotic, has shown the potential of OA disease amelioration by autophagy activation [17, 18, 27] and is proven to be senomorphic (senescence preventive drug) [30, 31]. However, the drawback of the free rapamycin use is low bioavailability at the target site even after local intra-articular injections. One of the strategies for extending the intra-articular half-life of therapeutics is encapsulation in polymer matrices which degrade slowly and release the drug over some time.

**Figure 7:**
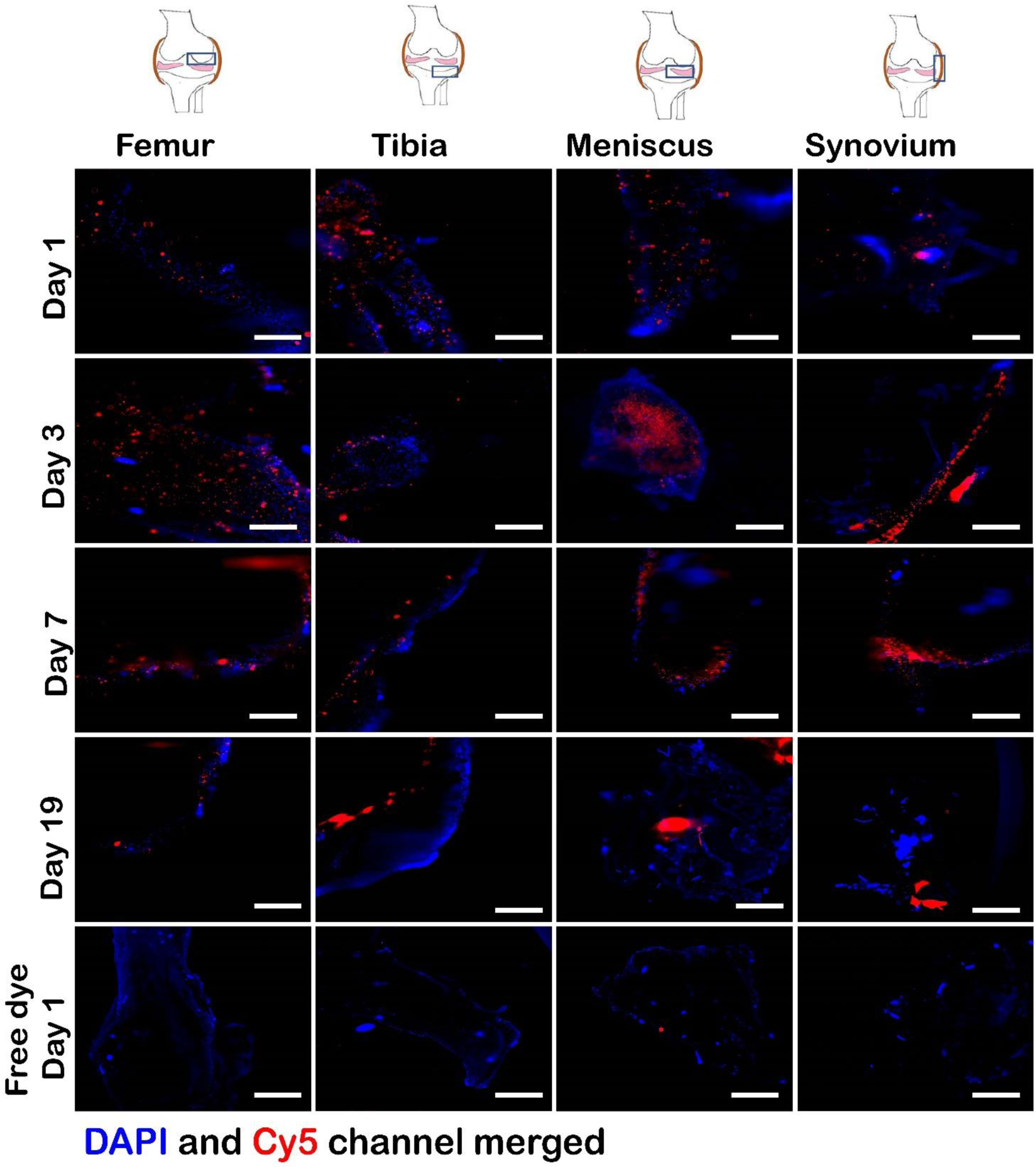
Cy5 loaded PLGA MPs of high molecular weight exhibit long residence time in mice knee joints. Fluorescence microscopy images of mice joints stained with DAPI and imaged at (a) 1 day, (b) 3 days, (c) 7 days and (d) 19 days after intra-articular injection of Cy5 labeled PLGA MPs. Rightmost insets depict the area of the joint that was imaged. **DAPI**-cells on the joint surface, **Cy5**-PLGA MPs encapsulating Cy5. Scale bar 100 µm.

Thus, using rapamycin and PLGA MPs as the drug carrier, we have developed a drug delivery system that provides the combined benefits of sustained local drug administration as well as prevents chondrocyte damage. Rapamycin was encapsulated in a microparticle formulation and stayed active and induced autophagy in chondrocytes in a dose dependent manner. The LC3B punctate signals in the rapamycin MPs treated group, indicative of autophagy activation, was comparable to that of the free rapamycin treated groups at the same dose.

It has also been reported that autophagy is essential to maintain the homeostasis in chondrocytes and downregulation of autophagy in cells pushes their entry into senescence while restoration of autophagy helps to reverse senescence [57, 58]. Senescence is a common molecular mechanism that drives or promotes both age-associated OA and trauma associated OA [8, 22]. Senescence occurs with prolonged multiplication of cells (replicative senescence) leading to aging and when stresses like nutrient deprivation or trauma occur to the cells (premature senescence) [54, 59, 60]. Once cells become senescent, they remain metabolically active and are known to induce senescence in surrounding cells and tissues [54].

Most of the current chondrosenescence research focuses on the selective killing of senescent in the OA milieu [23, 30, 31]. To the best of our knowledge, in this study, for the first time, we have successfully established rapamycin’s ability both as a free drug as well as a microparticle formulation to prevent the senescence in chondrocytes under genomic stress as well as oxidative stress conditions. The rapamycin MPs were able to prevent senescence in chondrocytes under stress in monolayer cultures. In 3D micro mass cultures, rapamycin MPs were able to sustain the production of sGAG, both after short and long duration of stress conditions. In cultures pre-exposed to oxidative stress for 2 days, rapamycin MPs were able to sustain sGAG production for up to 6 days even in the presence of continual stress. Prevention of cells from going into senescence and sustaining sGAG production can lead to enhanced cartilage repair and healing under stressed conditions like trauma or age induced OA. This can essentially pave the way for improving the joint health and can potentially slow down the onset of OA. Thus, a two-pronged approach of chondrocyte autophagy induction and prevention of chondrosenescence under stress led to sustained sGAG production using rapamycin MP formulation.

Getting therapeutics into joints in a targeted and sustained fashion is difficult, as soluble drugs exit the joints rapidly, via the capillaries (small molecules like drugs) and lymphatic system (macromolecules like proteins) [32-34]. The residence time reported range from as small as 0.23 h for acridine orange (M_w_ 370 Da) to 26.3 h for large molecules like hyaluronic acid [33]. These values exemplify the challenges faced by intra-articular therapy, especially for chronic conditions like OA. One of the strategies for extending the intra-articular half-lives of therapeutics includes the use of injectable polymer matrices which degrade slowly and release the drug over some time [32-34, 39]. The PLGA MPs formulation as a single intra-articular injection that we used here exhibited a residence time of longer than 19 days in the murine knee joints. Systems exhibiting such long residence time can be used to deliver rapamycin and other therapeutics for OA treatment with fewer injections making the system robust and safe. Given that both rapamycin and PLGA are already approved for human use, it could be rapidly translated. Further studies to test the therapeutic efficacy of rapamycin MPs in a preclinical model of OA are required to assess its translation potential in clinics.

## Conclusion

Treatment of osteoarthritis remains a challenge due to the absence of drugs that can prevent, reverse or halt the progression of the disease. A controlled drug release platform combined with the benefits of preventing the progression of OA disease can fill this need in clinics. The rapamycin MPs formulation developed in this study was capable of providing sustained release of the drug for over 20 days. Rapamycin MPs effectively induced autophagy on par with an equivalent dose of free rapamycin and prevented senescence in the human chondrocyte cell line. Further, treatment with rapamycin MPs maintained the sGAG production in 3D cultures after genotoxic and oxidative stress conditions. Finally, these particles had much higher joint residence (>19 days) over free dye after intra-articular injections. Such biomaterial-based delivery of rapamycin offers promise for further testing in animal models and clinical translation as a patient compliant treatment.

## Abbreviations

OA: Osteoarthritis,
WHO: World Health Organization,
DMM: Destabilization of Medial Menisci,
mTOR: Mammalian Target of Rapamycin,
IL-1RA: Interleukin -1 Receptor Agonist,
IA: Intra-articular,
MPs: Microparticles,
M_w_: Molecular weight,
PLGA: Poly (lactic-co-glycolic acid),
PLA: Poly Lactide,
PGA: Poly Glycolide,
KDa: Kilo Dalton,
DCM: Dichloromethane,
DMSO: Dimethyl Sulphoxide,
γH2Ax: Gamma Histone 2 Ax Protein,
FBS: fetal bovine serum,,
H_2_O_2_: hydrogen peroxide,
ROS: reactive oxygen species,
UV: ultraviolet,
MTT: 3-(4,5-dimethylthiazol-2-yl)-2,5-diphenyltetrazolium bromide,
CDS: Crystal dissolving solution,
Calcein AM: Acetoxymethyl,
PI: Propidium iodide,
DAPI: 4′,6-diamidino-2-phenylindole,
SA-β-Gal: Senescence Associated Beta Galactosidase,
sGAG: sulphated Glycosaminoglycans,
BrdU: 5-bromo-2’-deoxyuridine,
PFA: paraformaldehyde,
SEM: Scanning Electron Microscope,
FITC: Fluorescein isothiocyanate,

## Acknowledgements

We thank Dr Siddharth Jhunjhunwala and Dr Vaishnavi Ananthanarayanan for their valuable suggestions and Centre for BioSystems Science and Engineering (BSSE) for allowing us to access its facilities including GE Incell Analyzer and cell culture facilities. We also thank Dr Karthik Sunagar for use of Epoch Plate reader, Central animal facility (CAF) for breeding and maintaining mice, Molecular Reproduction Development and Genetics department (MRDG) central facilities for access to Tecan Infinite Pro ^200^ Plate reader. Advanced Facility for Microscopy and Microanalysis at Indian Institute of Science was used for SEM image acquisition.

## Author contributions

Kaamini M. Dhanabalan (KMD) and Rachit Agarwal (RA) designed research. KMD performed research on all experiments. Vishal K Gupta (VKG) performed SEM data acquisition. KMD and RA wrote the manuscript. All authors approved the final version to be published.

