## Supplementary Information for "Rapamycin-PLGA microspheres induce autophagy and prevent senescence in chondrocytes and exhibit long *in vivo* residence"

**Supplementary data**

Table S1: PLGA microspheres of different molecular weights and their respective sizes along with their encapsulation efficiency

| Molecular weight of PLGA  (Ratio of PLA: PGA) | Size in nm measured by DLS | Rapamycin encapsulation efficiency (%) |
| --- | --- | --- |
| 10KDa -14KDa (50:50) | 1021 ± 153 | 63.43 ± 1.326 |
| 24KDa-38KDa (50:50) | 1066 ± 174.9 | 48.91 ± 0.930 |
| 24KDa-38KDa (65:35) | 946 ± 155.9 | 41.42 ± 1.326 |
| 34KDa-54KDa (50:50) | 1070 ± 183.1 | 41.33 ± 8.040 |
| 75KDa-85KDa (50:50) | 1039 ± 188 | 32.6 ± 3.007 |

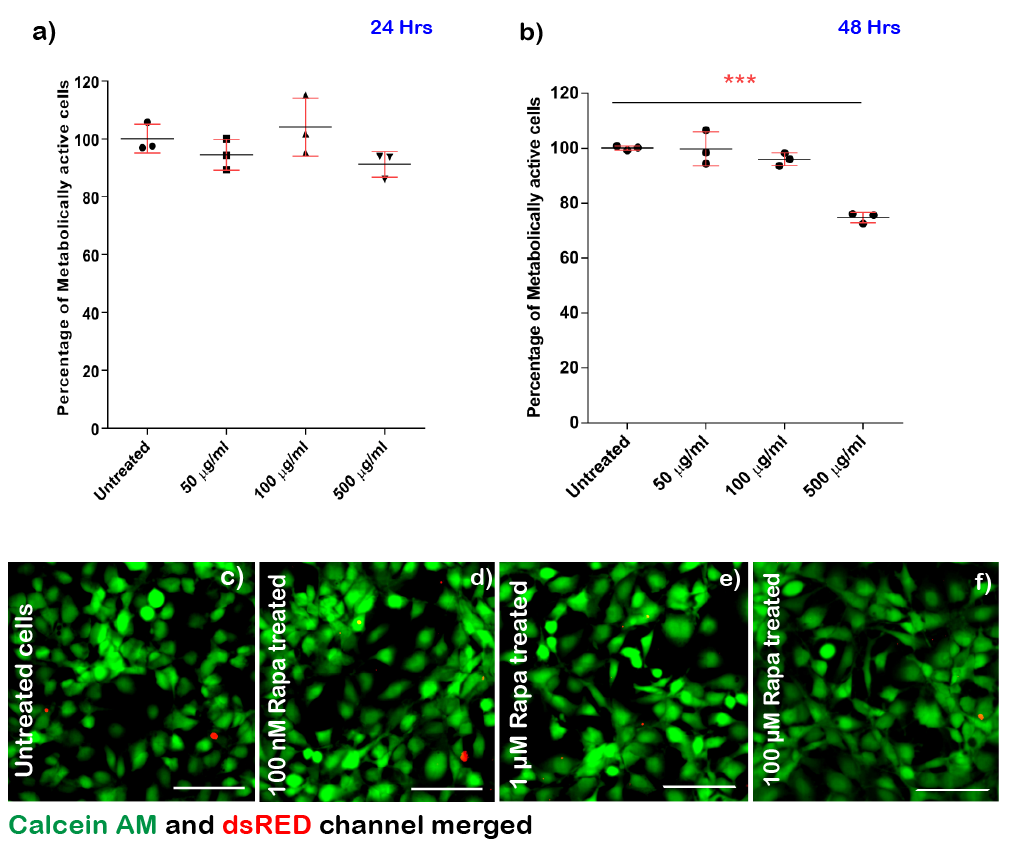

**Figure S1: Cytocompatibility assays for PLGA MPs and rapamycin.** Metabolic activity of chondrocytes treated with varying concentration of PLGA MPs for (a) 24 h and (b) 48 h. Images of Calcein AM (green) and Propidium Iodide (red) stained C28/I2 cells treated with (c) 0 nM, (d) 10 nM, (e) 1 μM and (f) 100 μM of Rapamycin. Scale bar 30 μm. ****p*< 0.001 for 500 μg/mL PLGA MPs versus all other groups. *p-*value < 0.05 were considered significant. Data in graphs represent the mean ± s.d. (n = 3) and *p* values were determined by One-way ANOVA and Tukey’s post hoc tests using GraphPad prism Software.

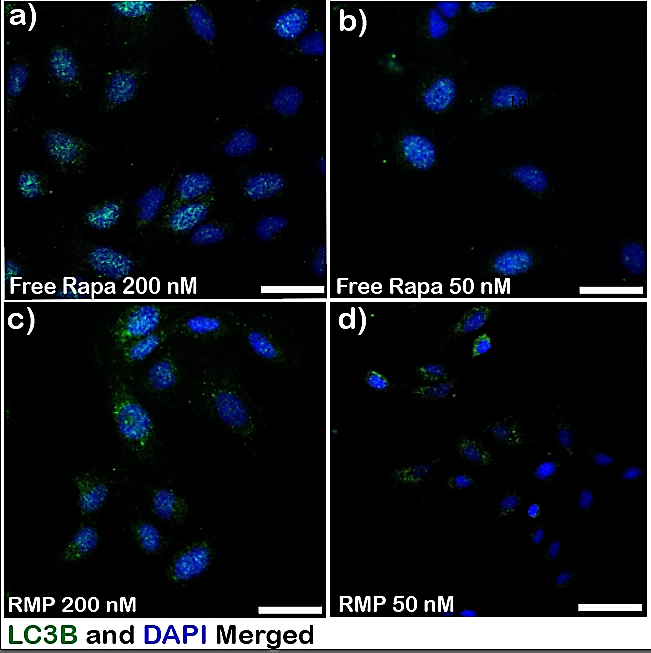

**Figure S2:** **Rapamycin MPs induces autophagy in chondrocytes.** Fluorescence microscopy images of C28/I2 cells stained with DAPI (blue) and LC3B (green) primary antibody after treatment with (a) Free Rapamycin 200 nM (b) Free rapamycin 50 nM (c) RMP (200 nM Rapamycin) (d) RMP (50 nM Rapamycin). Scale bar 20 μm.

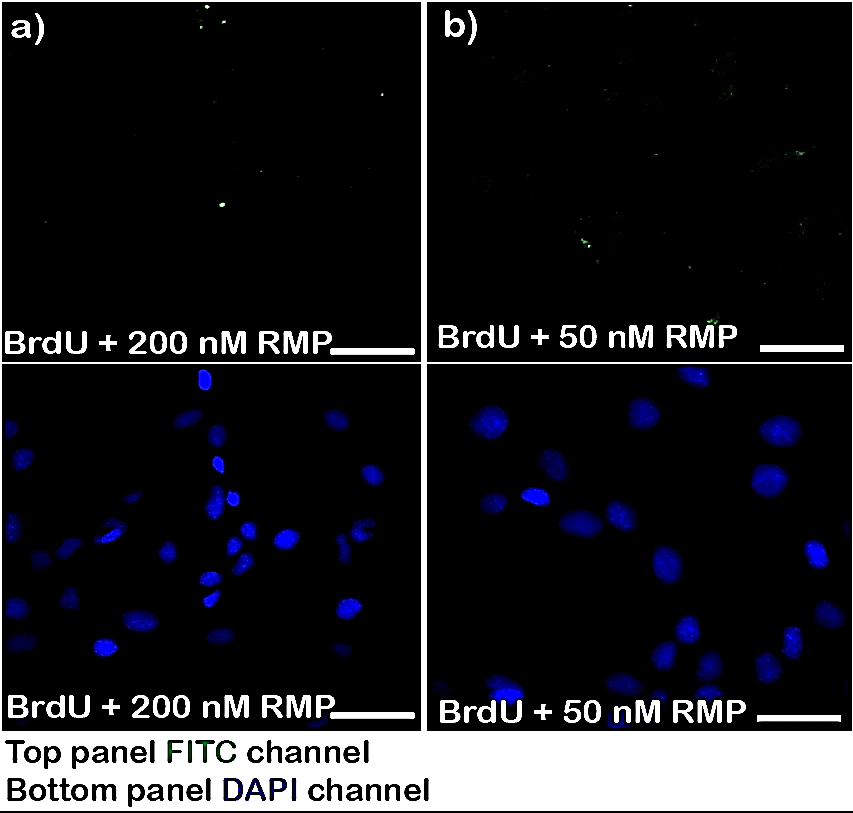

**Figure S3: Rapamycin MPs reduces DNA damage in chondrocytes.** Fluorescence microscopy images of C28/I2 cells stained with DAPI (blue) and γH2Ax (green) primary antibody after treatment with (a) BrdU (200 µM) along with Rapamycin MPs (200 nM) (b) BrdU (200 µM) along with Rapamycin MPs (50 nM). Scale bar 20 μm.
